# Does chronic systemic injection of the DREADD agonists clozapine-N-oxide or compound 21 change behavior relevant to locomotion, exploration, anxiety, and depression in male non-DREADD-expressing mice?

**DOI:** 10.1101/2020.05.17.100909

**Authors:** Fionya H. Tran, Stella L. Spears, Kyung J. Ahn, Amelia J. Eisch, Sanghee Yun

## Abstract

Designer Receptors Exclusively Activated by Designer Drugs (DREADDs) are chemogenetic tools commonly-used to manipulate brain activity. The most widely-used synthetic DREADD ligand, clozapine-N-oxide (CNO), is back-metabolized to clozapine which can itself activate endogenous receptors. Studies in non-DREADD-expressing rodents suggest CNO or a DREADD agonist that lacks active metabolites, such as Compound 21 (C21), change rodent behavior (e.g. decrease locomotion), but chronic injection of CNO does not change locomotion. However, it is unknown if chronic CNO changes behaviors relevant to locomotion, exploration, anxiety, and depression, or if chronic C21 changes any aspect of mouse behavior. Here non-DREADD-expressing mice received i.p. Vehicle (Veh), CNO, or C21 (1mg/kg) 5 days/week for 16 weeks and behaviors were assessed over time. Veh, CNO, and C21 mice had similar weight gain over the 16-week-experiment. During the 3rd injection week, CNO and C21 mice explored more than Veh mice in a novel context and had more open field center entries; however, groups were similar in other measures of locomotion and anxiety. During the 14th-16th injection weeks, Veh, CNO, and C21 mice had similar locomotion and anxiety-like behaviors. We interpret these data as showing chronic Veh, CNO, and C21 injections given to male non-DREADD-expressing mice largely lack behavioral effects. These data may be helpful for behavioral neuroscientists when study design requires repeated injection of these DREADD agonists.

**Highlights:**

- Acute injection of CNO changes behavior of non-DREADD-expressing mice
- It’s not known if chronic CNO or alternative agonist C21 also changes mouse behavior
- DREADD agonists or Veh were given chronically to non-DREADD-expressing mice
- CNO and C21 don’t change locomotion and have a mixed effect on anxiety-like behavior
- 1 mg/kg CNO and C21 can be injected repeatedly without non-specific behavior effects

## 1. Introduction

The preclinical use of chemogenetics, such as designer receptors exclusively activated by designer drugs (DREADDs), has enhanced manipulation of the brain activity in awake and behaving rodents [1–3]. The prototypical DREADD activator, Clozapine-N-oxide (CNO), was initially thought to be biologically-inactive [3]. However, CNO is now presumed to cause off-target endogenous receptor activation due to its back-metabolism to clozapine [4–7]. Clozapine is a clinical antipsychotic [8] and its acute administration to non-DREADD-expressing rodents decreases locomotion in a dose- and time-dependent manner [9–12] and can be either anxiolytic- [11–13] or anxiogenic-like [10,14] depending on dose. Thus, it is important to understand if the behavioral changes documented in studies that have used CNO are due to specific activation of DREADDs or to off-target effects via the non-DREADD-specific actions of clozapine. To avoid potential off-target effects, other designer drugs have been developed to interact with DREADDs [15,16], such as Compound 21 (C21) which has no back-metabolism to clozapine [16]. Notably, it is unknown if chronic administration of CNO or C21 to non-DREADD-expressing rodents changes behavior relevant to locomotion, exploration, anxiety, or depression. This is an important knowledge gap, as many studies administer DREADD agonists repeatedly [17,18]. Identifying the behavioral effect - or lack thereof - of chronic CNO or C21 in non-DREADD-expressing rodents would enable researchers to best adhere to the principles of the 3R’s (replacement, reduction, refinement) [19].

The behavioral relevance of back-metabolized clozapine from acute CNO remains unclear [9–14,20]. Some data support an off-target behavioral effect of acute CNO [5,7]; for example, non-DREADD-expressing mice given acute CNO (1mg/kg) locomote less than control mice when examined 2-3h post-injection, the predicted time point when back-metabolized clozapine concentration is highest [7]. However, other data do not support a behavioral or physiological effect of back-metabolized clozapine [6,21,22]; for example, in non-DREADD expressing animals, CNO (<5mg/kg) does not substitute for clozapine. To understand this apparent discrepancy - 1mg/kg acute CNO decreases locomotion, but <5mg/kg CNO is not discriminated - it is important to directly assess the influence of CNO on fundamental behaviors (e.g. locomotion) in non-DREAD-expressing rodents, and to examine these behaviors at the interval post-CNO when back-metabolized clozapine is thought to be highest. Also, since clozapine’s effect in both humans and mice is greater after chronic vs. acute administration [8,23], it is important to assess the behavioral effect of chronic CNO injections in non-DREADD-expressing rodents.

Here we assessed the behavioral effect of giving male mice chronic injections of Veh or the DREADD agonists CNO and C21. In line with best practices [5,6], non-DREADD-expressing mice were given Veh or a DREADD agonist at an experimentally-relevant dose (1mg/kg)[4,16] to discern off-target effects unrelated to DREADD activation. We hypothesized that relative to chronic Veh, chronic (like acute[6,7]) CNO would result in back-metabolized clozapine and thus decrease locomotion and exploration, while chronic C21 injections would result in similar performance in tasks relevant to locomotion, exploration, anxiety, and depression.

## 2. Materials and Methods

### 2.1 Ethics

Experiments were approved by the Institutional Animal Care and Use Committee at the Children’s Hospital of Philadelphia (CHOP) and performed in compliance with NIH’s *Guide for the Care and Use of Laboratory Animals*.

### 2.2 Animals and Genotyping

Fifteen-week-old, experimentally-naive and non-DREADD-expressing male *B6.FVB-Tg (Camk2a-cre) 2Gsc/Cnrm* mice (*CamKIIα-icre*, congenic C57BL/6J, n=24, MGI: 3694800 [18,24]) were bred, housed, and weighed weekly at CHOP’s AAALAC-accredited, specific-pathogen-free conventional vivarium **(Supplementary materials)**.

### 2.4 Drugs

Each cage (2-4 mice) was randomly-assigned to the Veh, CNO, or C21 group (n=8/group). At 15-weeks-of-age (**Fig. 1A**), mice began receiving daily (Monday-Friday, 11:00am-1:00pm) i.p. injections of Veh (5ml/kg 0.5% dimethyl sulfoxide, 0.9% Saline), CNO (NIMH Chemical Synthesis and Drug Supply Program; 0.2mg/ml in Veh, 5ml/kg for final 1mg/kg dose), or C21 (Hello Bio, #HB4888; 0.2mg/ml in 0.9% Saline, 5ml/kg for final 1mg/kg dose). Injections were given between 11:00am-1:00pm and in counterbalanced-order by cage 3h prior to most behavior tests, 90 minutes (min) prior to locomotion testing on Day1, and 60min into activity monitoring.

**Figure 1.**
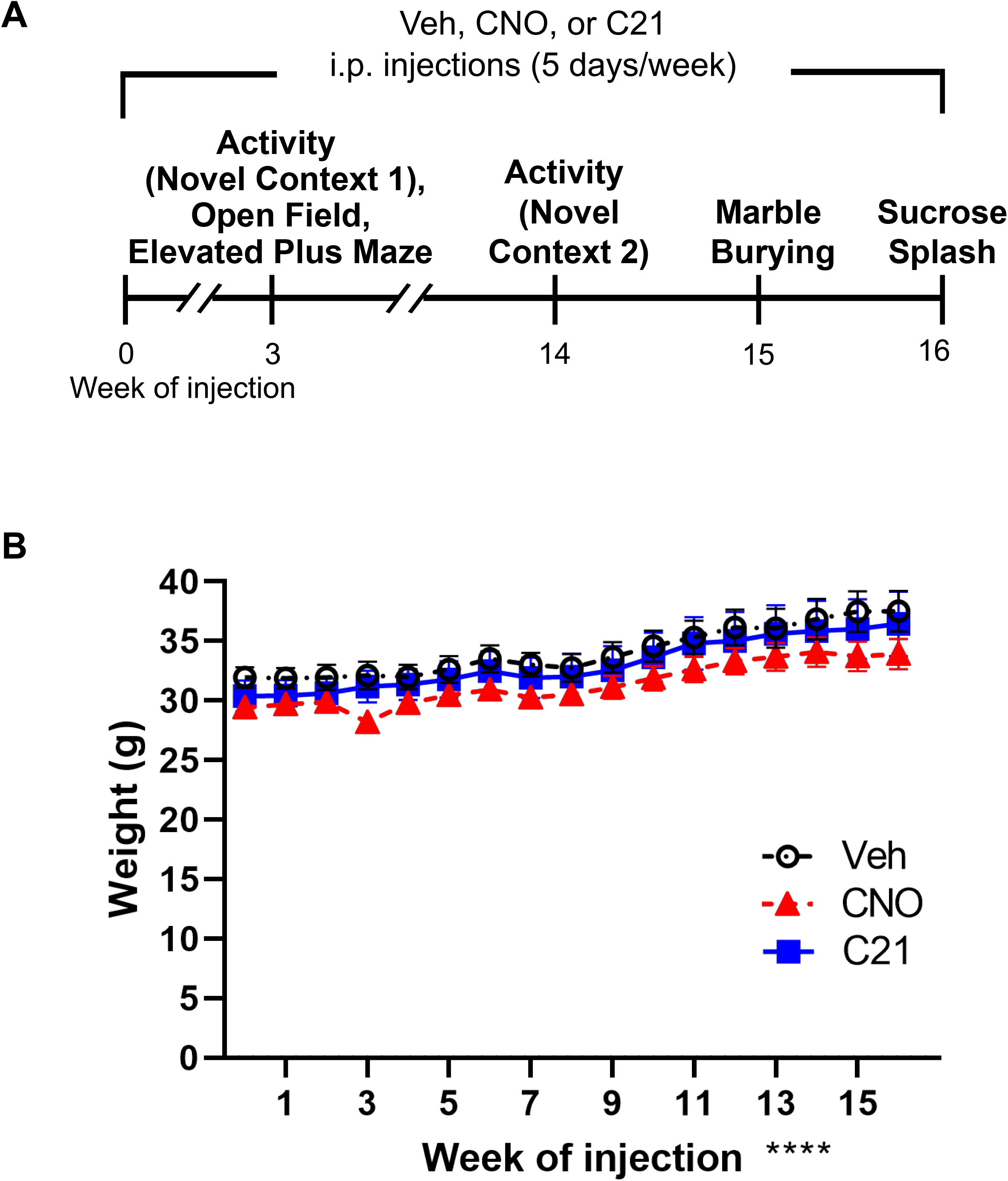
Timeline of behavior tests and body weight data. **(A)** Behavioral tests began after three weeks of i.p. injections. **(B)** Mouse weight gain did not differ by Treatment but did differ by Week.

### 2.5 Behavior

Behavior testing proceeded as shown (**Fig. 1A**). Since clozapine back-metabolized from CNO peaks 2-3h after CNO injection, and behavior changes 3-4h post-injection [6,7,18,25], most behaviors were tested 3h post-injection.

### 2.6 Statistics

Main effects and interactions were considered significant at P<0.05, and Bonferroni post-hoc tests were then performed. Results and **Supplementary Table1** provide effect sizes (omega-squared, ω^2^; partial omega-squared, ω_p_^2^) and P-values to four significant digits. All data are presented in the Results, but conclusions are not stated when subject numbers fell below the predetermined threshold **(Supplementary materials)**.

## 3. Results

### 3.1. Body weight, Weeks 0-16

Veh, CNO, and C21 groups gained a similar amount of weight at a similar rate (**Fig. 1B**; Mixed Measure Analysis, main effects of Treatment (F_2,21_=1.036, P=0.3722, ω^2^ =0.00) and Week (F_16, 326_=51.77, ****P<0.0001), and Interaction of TreatmentXWeek F_32,326_=0.6287, P=0.9438).

### 3.2 Activity (Novel Context 1), Week 3

The measure of Day1 exploration **(Fig. 2A,C)** showed threshold main effects of Treatment in both total beam breaks (F_2,21_=2.949, P=0.0744, ω^2^ =0.14, “large”) and 5min-bin breaks (F_2, 21_=3.35, P=0.0546, ω_p_^2^ =0.25, “large”) and a main effect of Time (F_3.755, 78.84_=13.05,****P<0.0001, ω_p_^2^ =0.42, “large”); Subject F_21, 105_=5.228,****P<0.0001) but no TimeXTreatment interaction (F_10, 105_=1.43, P=0.1995, ω_p_^2^ =0.02). Day2 exploration (**Fig. 2E,G**) showed a main effect of Time (F_3.788, 79.54_=13.62,****P<0.0001, ω_p_^2^ =0.21, “large”) not Treatment (total:F_2,21_=1.29, P=0.2962, ω^2^ =0.14, “large”; 5min-bins:F_2, 21_=1.63, P=0.2198, ω_p_^2^ =0.07; Subject F_21, 105_=5.917,****P<0.0001) and a TimeXTreatment interaction (F_10, 105_=1.928, P=0.0492; post-hoc Ps>0.122, ω_p_^2^ =0.04). The measure of Day1 locomotion (**Fig. 2B,D**) showed a main effect of Time (F_2.692, 56.53_=31.05,****P<0.0001, ω_p_^2^ =0.47, “large”) not Treatment (total:F_2, 21_=2.132, P=0.1436, ω_p_^2^=0.06; 5min-bins:F_2, 21_=2.132, P=0.1436, ω_p_^2^ =0.09; Subject F_21, 105_=2.974,****P=0.0001), and a TimeXTreatment interaction (F_10,105_=1.670, P=0.0974, ω_p_^2^=0.04). Day2 locomotion (**Fig. 2F,H)** showed a main effect of Time (F_3.725, 78.23_=3.057, P=0.0241, ω_p_^2^=0.05; Subject F_21, 105_=3.381,****P<0.0001) and a threshold TimeXTreatment interaction (F_10,105_=1.866, P=0.0582, ω_p_^2^=0.04) but no Treatment effect (total:F_2, 21_=0.0239, P=0.9764, ω_p_^2^=0.09; 5min-bins:F_2, 21_=0.1584, P=0.8545, ω_p_^2^=-0.01). Thus during the 3rd injection week in Activity Day1, CNO and C21 increased exploration but not locomotion.

**Figure 2.**
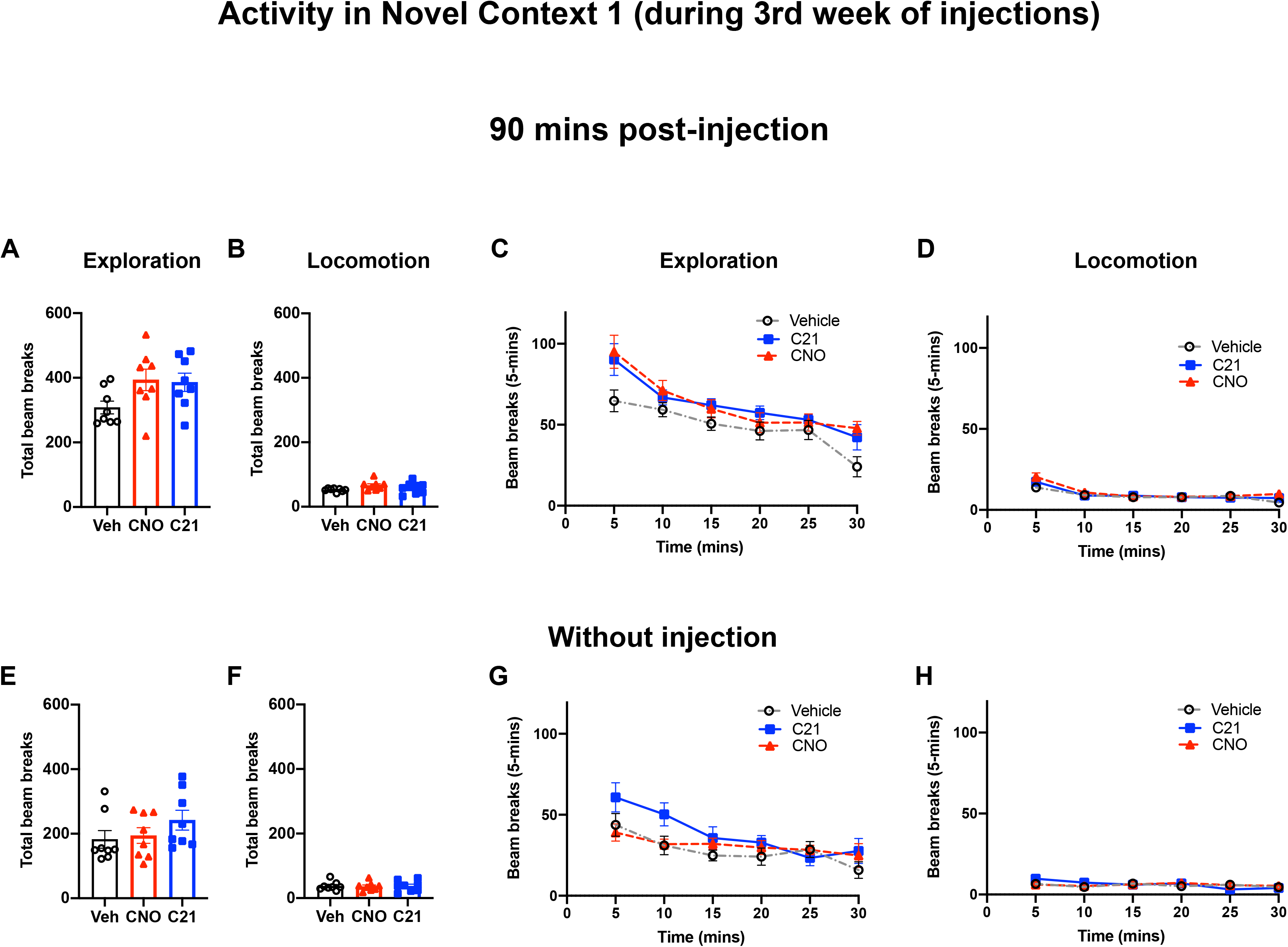
Activity (Novel Context 1) after three weeks of injection of Veh, CNO, and C21: exploration and locomotion. **(A-D)** Day1: Activity in novel context 1 90 min after injection of Veh, CNO, or C21 as measured by indices of exploration **(A, C)** and locomotion **(B, D)** on total beam breaks **(A-B)** and beam breaks/5min **(C-D)**. **(E-H)** Day 2: 24h after most recent injection of Veh, CNO, and C21, activity as measured by indices of exploration **(H, I)** and locomotion **(J, K)** on total beam breaks **(H-F)** or beam breaks/5min **(G-H)**. Mean±SEM. n=8/group.

### 3.3 Anxiety-relevant tests, Week 3

In the open field (**Fig. 3A-C**), Veh, CNO, and C21 mice moved a similar total distance (F_2, 21_=0.5440, P=0.5884, ω^2^ =-0.92). There was a main effect of Treatment on center entries (F_2, 21_=3.636, P=0.0441, ω^2^ =0.18, “large”) but no post-hoc significance. There was a threshold main effect on center zone duration (F_2, 21_=3.308, P=0.0564, ω^2^ =0.06). In the elevated plus maze (**Fig. 3D-F**), Veh, CNO, and C21 mice had similar total distance moved (F_2, 21_=0.3409, P=0.715, ω^2^ =-0.06) and open arm entries (F_2, 21_=0.9192, P=0.414, ω^2^ =-0.01) and duration (F_2, 21_=1.235, P=0.311, ω^2^ =-0.02). Thus, during the 3rd injection week, CNO and C21 have mixed effects, lowering anxiety/raising exploration in the open field but not changing elevated plus maze behavior.

**Figure 3.**
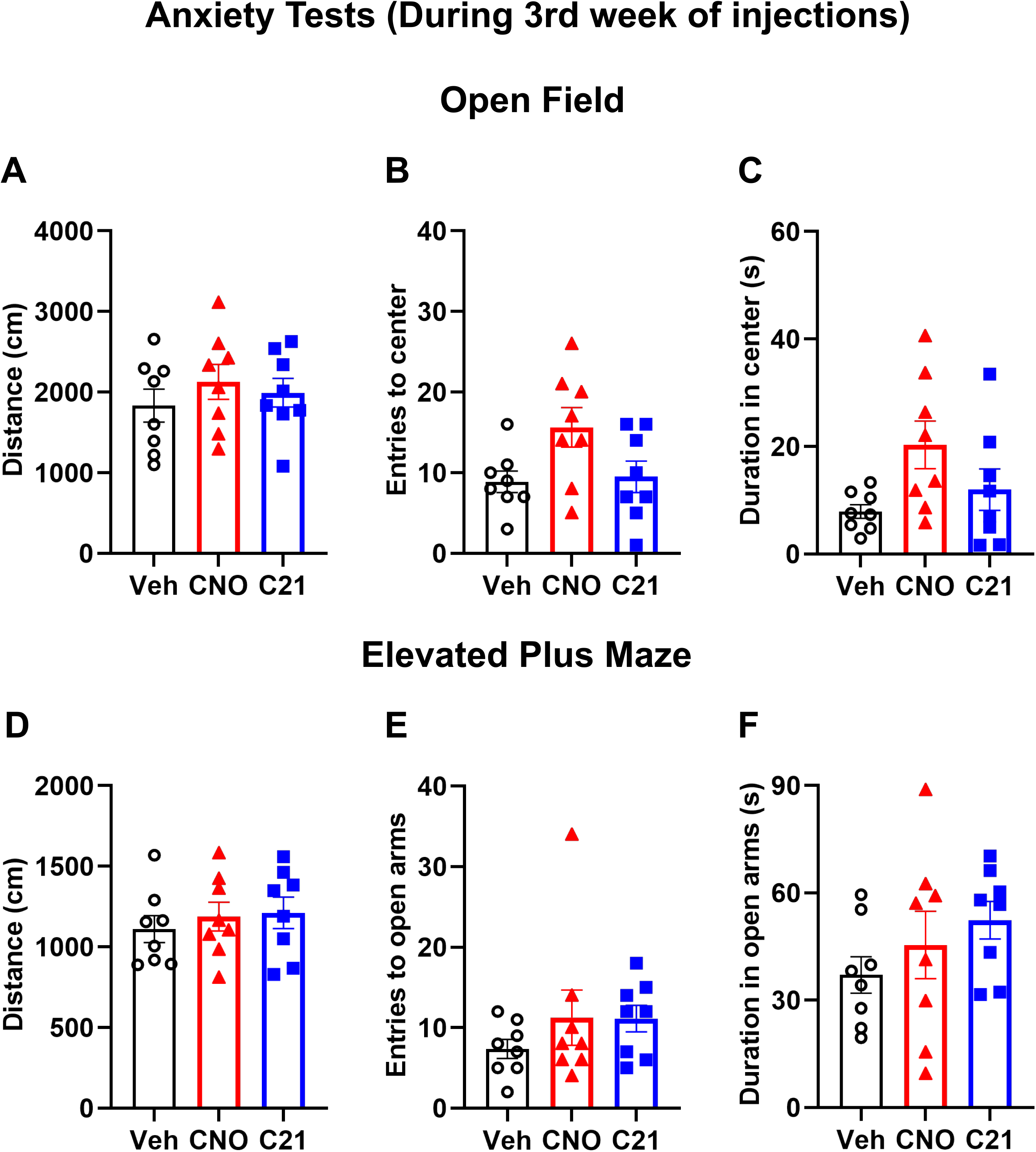
Open field and elevated plus maze after three weeks of injections of Veh, CNO, or C21: exploration, locomotion, and anxiety-like behaviors. **(A-C)** Mice given Veh, CNO or C21 had similar performance in the open field, as based on measures of total distance moved **(A)**, entries into the center zone **(B)**, and time spent in the center zone **(C)**. **(D-F)** Mice given Veh, CNO, or C21 had similar performance in the elevated plus maze, as based on measures of total distance moved **(D)**, entries into the open arms **(E)**, and time spent in the open arms **(F)**. Mean±SEM. n=8/group.

### 3.4 Activity (Novel Context 2), Week 14, and after acute CNO

Veh, CNO, or C21, ambulatory distance, duration, and movement events (**Fig. 4A-C**) showed a main effect of Time (distance:F_7, 140_=68.45,****P<0.0001,ω_p_^2^=0.68, subject F_20, 140_=3.597,****P<0.0001; duration:F_7, 140_=61.56,****P<0.0001,ω_p_^2^=0.7; subject:F_20, 140_=3.429,****P<0.0001; movement:F_7, 140_=61.56,****P<0.0001,ω_p_^2^=1.32; Subject F_20, 140_=3.429,****P<0.0001), a threshold effect of Treatment only in distance (F_2, 20_=3.014, P=0.0718, ω_p_^2^=0.08) but not in duration (F_2, 20_=2.298, P=0.1263, ω_p_^2^=0.06) or movement (F_2, 20_=1.900, P=0.1756, ω_p_^2^=0.05), and no TimeXTreatment interaction (distance:F_14, 140_=0.5073, P=0.507,ω_p_^2^=0.00; duration:F_14, 140_=0.8441, P=0.62,ω_p_^2^=0.00; movement:F_14, 140_=0.8441, P=0.62,ω_p_^2^=-0.01). Total jumps did not show a Treatment effect (F_2, 20_=0.5659, P=0.576, ω_p_^2^=-0.04). Thus, during the 14^th^ injection week, Veh, CNO, and C21 result in similar activity. To complement these chronic data, we assessed if acute 1mg/kg CNO changed activity in non-DREADD-expressing mice as reported by some [7,26] but not others [22]. Naive mice given a single injection of Veh or 0.3 or 1mg/kg CNO appeared to have similar activity over the next 16h **(Supplementary Fig. 1)**, but subject numbers were too low to state a conclusion.

**Figure 4.**
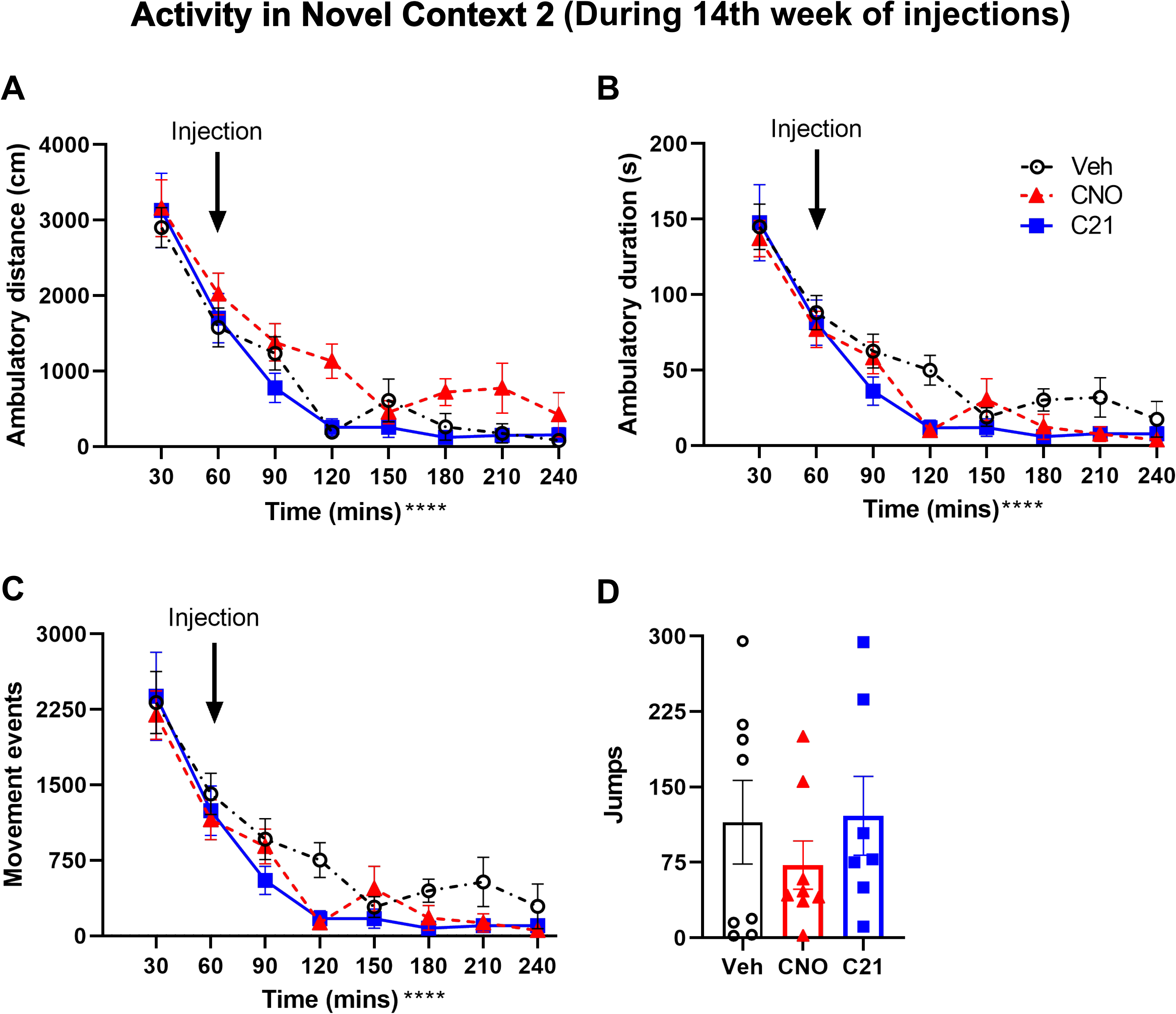
Activity (novel context 2) after fourteen weeks of injections of Veh, CNO, or C21: exploration and locomotion. **(A-D)** Mice given Veh, CNO, or C21 had similar activity in a novel environment based on ambulatory distance **(A)**, ambulation duration **(B)**, movement events **(C)**, and jumps **(D)**. Mean±SEM. Veh and CNO n=8, C21 n=7.

### 3.5 Tests relevant to anxiety and depression, Weeks 15-16

In marble burying (**Supplementary Fig. 2A**), Veh, CNO, and C21 mice buried a similar percentage of marbles (F_2, 20_=0.09887, P=0.906, ω_p_^2^=-0.09), indicating similar behavior relevant to anxiety and/or repetitive action. In the sucrose splash test (**Supplementary Fig. 2B-C**), data suggest Veh, CNO, and C21 mice had similar measures on latency to grooming and grooming events; however, the Veh subject number was too low to state a conclusion.

## 4. Discussion

Here we examined the behavioral effects of chronic CNO or C21 injections in non-DREADD-expressing mice. We administered 1mg/kg based on the ability of this dose to change behavior in DREADD-expressing rodents when injected acutely [16,27] or chronically [18,28]. We assayed most behaviors 3h post-injection, when CNO-to-clozapine back-metabolism peaks [4,7], and tested locomotor activity 0-3h post-injection. For the non-DREADD-expressing mouse line, we selected *CamKIIa-icre* mice given the use of this forebrain glutamatergic neuron cre-expressing line in DREADD studies by us and others [1,18,24,29]. Data from the 3rd injection week show these non-DREADD-expressing mice given chronic CNO or C21 explored a novel context more (but had similar locomotion) relative to mice given Veh, and showed mixed responses in anxiety tests. In all other tests during the 16-week experiment, Veh, CNO, and C21 mice had similar behavior; the effect of chronic CNO or C21 on depressive-like behavior (splash test) and acute CNO on locomotion was inconclusive due to loss of subjects. Overall, our data suggest that when appropriate control groups are used, non-DREADD-expressing mice can be injected repeatedly with either DREADD agonist without causing gross behavioral effects.

Our largely-negative data are consistent with reports that chronic 1mg/kg CNO given to non-DREADD-expressing mice does not change behavior/physiology [28,30], but our data importantly show no effect on locomotion and a mixed effect on anxiety-like behavior. While the literature is unanimous that chronic 1mg/kg CNO does not grossly change mouse behavior in non-DREADD-expressing mice, the literature is mixed on the effects of acute CNO given to non-DREADD-expressing rodents. Several studies report acute CNO or C21 does not change behavior in non-DREADD-expressing rodents (1mg/kg [our data, 22] or 3 mg/kg [4,21] CNO, or 3.5mg/kg C21 [4]). Of studies that do report a change in behavior in non-DREADD-expressing rodents after acute CNO, two saw behavioral effects in rats with acute 1mg/kg CNO [7,26] and two in mice and rats with higher CNO doses than the present work [4,6]. Further work is needed to understand what factors (mouse strain, behavior parameters, etc.) underlie these divergent results with acute CNO.

CNO and C21 are both rapidly-metabolized in mice [4], but brain C21 levels last slightly longer than brain CNO levels [4]. Thus, C21 may be a better designer drug for studies requiring longer duration of DREADD activation. While C21 binds to a range of G-protein coupled receptors (including dopamine D1 and D2 and histamine H4 receptors) at doses >3mg/kg, below this dose it appears to be a reliable activator since unlike CNO it has no detectable conversion to either clozapine or CNO [4,16].

## Conclusion

While Non-DREADD-expressing mice given CNO or C21 (1mg/kg i.p) for 5 days/week for 3-16 weeks perform indistinguishably from Veh mice in tests relevant to locomotion, mice given CNO or C21 have increased exploration during the 3rd injection week; results on anxiety measures were mixed. These data suggest with appropriate dose and control groups, CNO and C21 can be used as DREADD agonists for studies that require long-term, repeated injection of these compounds without concern for gross non-specific behavioral or physiological effects.

## Declaration of Competing Interests

Authors have no conflicts of interest to declare.

## Acknowledgements

This work was supported by the University of Pennsylvania McCabe Fund Pilot Award (SY), a 2019 International Brain Research Organization travel grant (SY), the 2019 Brain & Behavior Research Foundation NARSAD Young Investigator Grant (SY), University of Pennsylvania Undergraduate Research Foundation grant (SY), the National Aeronautics and Space Administration (AJE, 80NSSC17K0060), the Children’s Hospital of Philadelphia Department of Anesthesiology and Critical Care Development Funds (AJE), and the University of Pennsylvania Office of the Vice Provost for Research (SLS). We also acknowledge the generous intellectual engagement and support of the members of the Eisch Lab.

## Abbreviations

CNO: clozapine-n-oxide
C21: compound 21
DREADD: designer receptor exclusively activated by designer drugs
h: hour
i.p.: intraperitoneal
min: minutes
Veh: vehicle

## Supplementary materials

### S1. Materials and Methods

#### S1.1 Animals and Genotyping

*CamKIIα-iCre* hemizygous mice were bred from homozygous *CamKIIα-iCre* male mice crossed with C57BL/6J female mice. Mice received ear tags at four weeks of age (National Band and Tag Company, #1005-1L1). Mice were kept on a 12-h light/dark cycle (lights on 6:15am) with *ad libitum* food (Lab Diets 5015 #0001328) and water. Room environments were maintained at 20-23°C and 30-70% humidity. Home cages consisted of individually-ventilated polycarbonate microisolator cages (Lab Products Inc., Enviro-Gard™ III, Seaford, DE) with HEPA filtered air, corncob (Bed-o’ Cobs^®^ ¼”) bedding, one nestlet (Ancare), and a red plastic hut (Bio-Serv, #K3583 Safe Harbor). Mouse genotype (hemizygous CamKIIα-icre males) was verified by following previously published methods [1]. Genomic DNA is extracted from the ear punch samples. Ear samples were heated in 200ul 0.05M NaOH solution at 95°C for 10min and were neutralized with 50ul 0.5M TRIS HCl, pH 8.0 following centrifugation (10,000 rcf for 10min). The supernatant underwent PCR using primers for Cre recombinase (TGG TGC CCA AGA AGA AGA GGA A; CAT TCT TTC TGA TTC TCC TCA TCA) and GoTaq^®^ Green Master Mix (Promega, M7123). An amplified DNA band for cre recombinase was identified by gel electrophoresis with EtBr.

#### S1.2 Drug

Chronic injection procedure is provided in the main text. For the acute injection experiments, mice began receiving i.p. injections of Veh (5ml/kg 0.5% dimethyl sulfoxide, 0.9% Saline) or CNO (NIMH Chemical Synthesis and Drug Supply Program; 0.2mg/ml in Veh, 5ml/kg for final 0.3 or 1mg/kg dose) right before activity monitoring.

#### S1.3 Weight (0-16th injection week)

Mice were weighed every Monday using an OHAUS™ Valor^®^ 2000 scale (#V22PWE1501T). Due to building construction, mice were moved to a different CHOP animal facility during the 7th injection week for the study remainder. After this move, one C21 mouse had unexplained weight loss and was therefore removed from the study.

#### S1.4 Overview of behavior testing for mice in the chronic injection study

Testing in mice given chronic injection of Veh, CNO, or C21 proceeded as noted in **Fig. 1A**. In the 3rd injection week, mice underwent activity monitoring in novel context 1 (2 consecutive days), open field [1], and elevated plus maze [1]. In the 14th injection week, mice underwent activity monitoring in novel context 2. In the 15th and 16th injection weeks, mice underwent marble burying [2] and sucrose splash test [1], respectively. Initial study design included the forced-swim test, but as three Veh mice died during initial forced-swim testing, no other mice were run on this test. Prior to each test, mice were habituated to the testing room for ~1h under red light (30-50 lux).

##### S1.4.1 Activity in Novel Context 1 (3rd injection week)

Activity in a novel context (termed novel context 1) was tested 30min/day for two consecutive days (Day 1: CNO given 90 min prior to testing, Day 2: no CNO given). Mice were individually placed into a thoroughly-cleaned trapezoidal Bussey-Saksida Touch System Chamber (Lafayette Instruments, #80614) with the widest wall serving as the “touchscreen” (W23.8xH17 cm) and the opposite and narrowest wall (W4.6cm) containing a motion-sensitive center tray. The center tray remained empty and no images were presented on the touchscreen wall during the 30min test. The test was performed at lux 0. The chamber had infrared detection beams running across the center tray, directly in front of the wall opposite the center tray (the touchscreen), and across the chamber width (in parallel with infrared beams across center tray, termed chamber activity beams). Beam breaks in 5min-bins and total beam breaks over the 30min test were recorded by ABET II software (Lafayette Instruments). Measures relevant to exploration vs. locomotion were analyzed separately. Exploration was defined as vertical and horizontal movements within the center tray or on the opposite wall. Locomotion was defined as total or partial movement detected by chamber activity beams (movement between wall and reward hopper).

##### S1.4.2 Open Field (3rd injection week)

Three hours after injection, mice underwent open field testing [1] during the 3rd injection week. The test was performed at 35-50 lux. Mice were placed in a square (L42xW42xH42cm) opaque-white Plexiglas chamber (Nationwide Plastics). A single mouse was allowed 5-min of free movement in the chamber. Total movement distance, center zone (L14xW14cm) entries and duration, and peripheral zone (L5xW5cm) entries and duration were scored via EthoVisionXT software (Noldus Information Technology) using nose-center-tail tracking.

##### S1.4.3 Elevated Plus Maze (3rd injection week)

Three hours after injection, mice were placed in the elevated plus maze apparatus [1] (black Plexiglas floor, Harvard Apparatus, #760075) had two open arms (L67xW6cm) and two closed arms with walls (L67xW6xH17cm, opaque-grey Plexiglas walls). The test was performed at 35-50 lux. Mice were placed at the far end of one open arm and allowed 5-min free movement. Total distance moved, open arm entries and duration, and closed arm entries and duration were scored via EthoVisionXT software (Noldus Information Technology) using nose-center-tail tracking.

##### S1.4.4 Activity in Novel Context 2 (14th injection week)

Square chambers (L27xW27xH20cm, Med Associates Inc., #ENV-510), each placed inside an opaque sound-attenuating chamber, were used to measure activity via infrared beam breaks over 240min.The test was performed at lux 0. After 60min, mice were removed from the square chambers, injected (Veh, CNO, or C21), and immediately placed back into the chamber for the final 180min. Measures of total ambulatory distance, ambulatory duration, number of movement events, and total number of jumps were scored with Activity Monitoring 5 software (Med Associates Inc., #SOF-811). Each measure was analyzed separately.

##### S1.4.5 Marble Burying (15th injection week)

Three hours after injection, marble burying [2] was performed in a fresh transparent polycarbonate cage (L25.7xW48.3xH15.2cm, with filter-top lids; Allentown Inc. #PC10196HT) with 5cm of Beta Chip Bedding (Animal Specialties and Provisions, #NOR301) covering the cage bottom. The test was performed at 35-50 lux. Twenty glass marbles (13mm diameter, cat’s eye design, primary colors) were placed on the bedding (4×5rows). Mice were placed in the cage for 20min. The number of marbles buried (covered ⅔ or more in bedding) by the end of the test was scored by two independent observers and averaged.

##### S1.4.6 Sucrose Splash Test (16th injection week)

The day before the sucrose splash test, mice were singly-housed in fresh microisolator cages. The next day mice were given their respective injection, and the nestlet, food hopper, and red plastic hut were removed from the home cage right injection. For the test itself (performed at 35-50 lux), freshly-made 10% sucrose in water was sprayed onto the mouse’s back [1]. Video recordings (5min, Panasonic, HC-V270) were manually scored by an observer unaware of treatment group. Measures reported are total grooming duration, latency to groom, and total grooming events.

#### S1.5 Activity in Home Cage after Acute Veh or CNO

Activity monitoring in a home cage **(Supplementary Fig. 1)** was measured as previously described [3]. In brief, mice received a single i.p. injection of Veh, 0.3mg/kg CNO, or 1 mg/kg CNO and were immediately and individually placed in a fresh home cage (same dimensions as housing cage, but fresh bedding, food, and water) between photocells. The test was run under dim light (30-50 lux; 5pm-6:15pm, 6:15pm-9:00am) and red light (30 lux; 6:15pm-6:15am). A computer-controlled photobeam activity system (San Diego Instruments) recorded total movement of mice in the XY plane, with photocell beam breaks recorded with 15min-bins overnight (16h, from 17:00 to 09:00).

#### S1.6 Rigor, statistical analyses, and effect size determination

Experimenters were blinded to treatment until behavior was complete. Data are reported as mean±SEM and were analyzed as noted in **Supplementary Table 1**. All data were analyzed via null-hypothesis significance testing (NHST; Prism 8 GraphPad Software, San Diego, CA) and with estimation statistics (www.aggieerin.com/shiny-server/tests/omegaprmss.html) per best practice recommendations [4–7]. For NHST, one-way ANOVA was performed for activity, open field, elevated plus maze, marble burying, and sucrose splash test. A repeated measure (RM, within-subject time) two-way ANOVA was performed for activity beam breaks over time. Due to varied group size, body weight and activity were analyzed via mixed measure (within-subject repeated measure, between-subject non-repeated). Bonferroni post-hoc tests were performed when P-values<0.05 for the main effect of Treatment or when there was an interaction between two independent variables. In this work, the word “threshold” is applied to a main effect when 0.05<P<0.1, and confidence intervals (CI, 95%) for one-way ANOVAs are provided in **Supplementary Table 1**. For estimation statistics, omega-squared (ω^2^) was used for one-way ANOVA and partial omega-squared (ω_p_^2^) was used for RM twoway ANOVA. In line with the literature [4–7], ω^2^ and ω_p_^2^ effect sizes were considered as following: ≤0, similar to 0; 0.1, small; 0.06, medium; and 0.13, large. All effect sizes (ω^2^ and ω_p_^2^) are provided in the Results and **Supplementary Table 1** regardless of NHST P value, and the classification of ω^2^ and ω_p_^2^ effect sizes are only provided in the Results for “large” effect sizes. All behavioral data collected were analyzed and are presented in the Results. However, conclusions are not stated for the splash test or acute injection activity as these subject numbers fell below the predetermined threshold.

**Supplementary Table 1.**
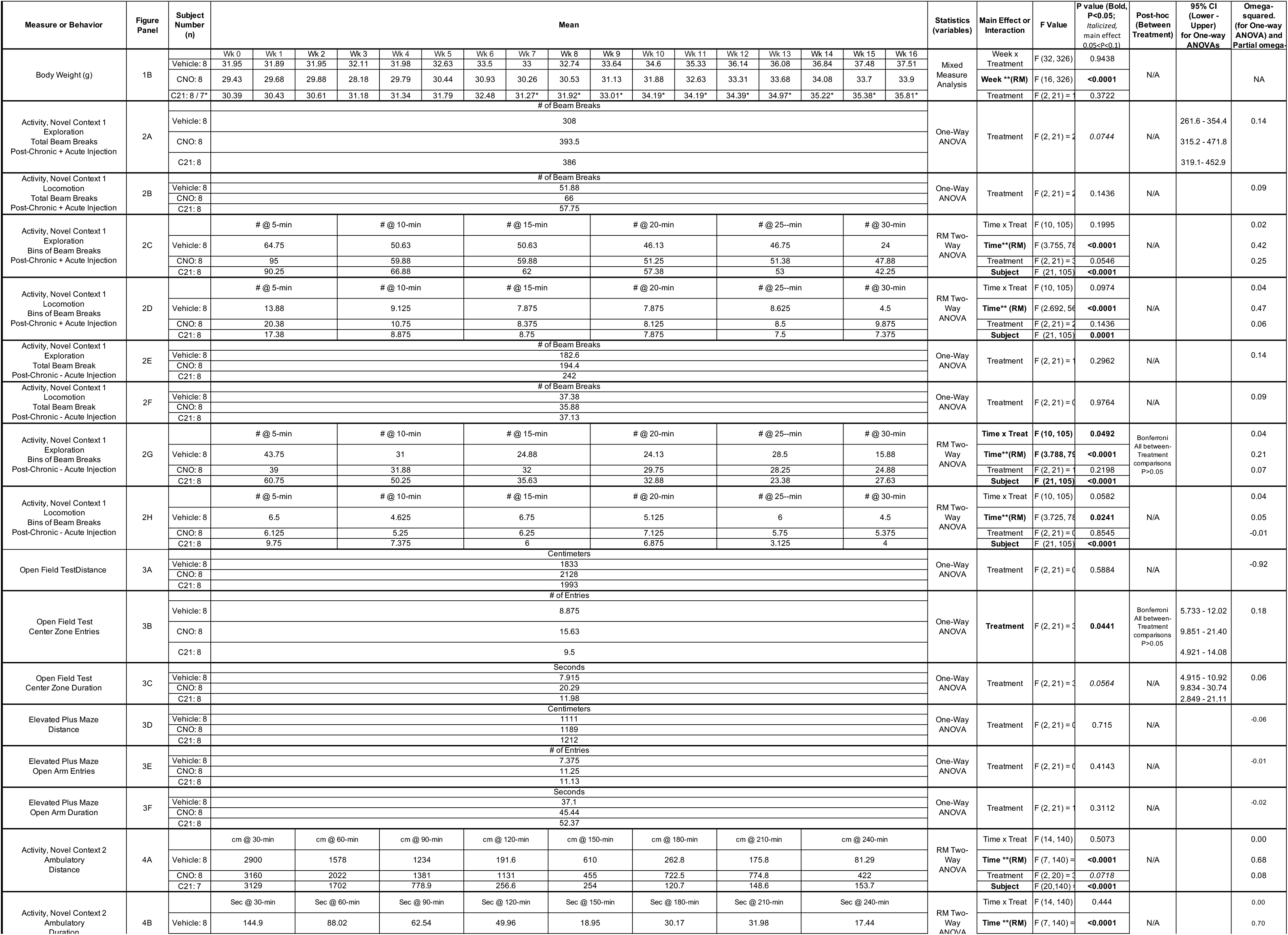
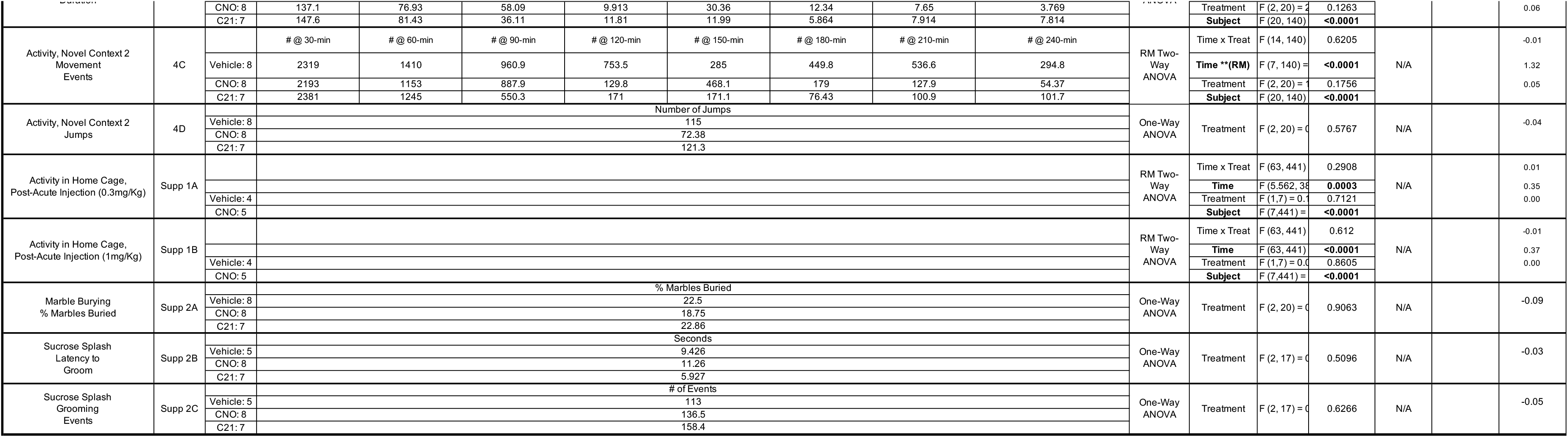
Statistical details for each main and supplementary figure panel provided in Tran, Spears, et al. As in the Results, P values are provided up to four significant digits. **Bold text**, P<0.05. *Italicized text* is for “threshold” effects where 0.05<P<0.1. 95% confidence intervals (lower and upper) are provided for one-way ANOVA results when P<0.1. Omega squared (ω^2^; for one-way ANOVA) and partial omega squared (ω_p_^2^; for RM two-way ANOVA) effect sizes are provided for all results with the exception of weight data, as it could not be calculated due to missing values. Effect size based on values of ω^2^ and ωp: 0.01 small; 0.06 medium; 0.13 large. * loss of C21 mouse during Week 7. ** Factors repeated in Mixed Measure Analysis. N/A Not applicable (no main effect of Treatment, so no post hoc test performed).

### Supplementary Figure Legends

**Supplementary Figure 1.**
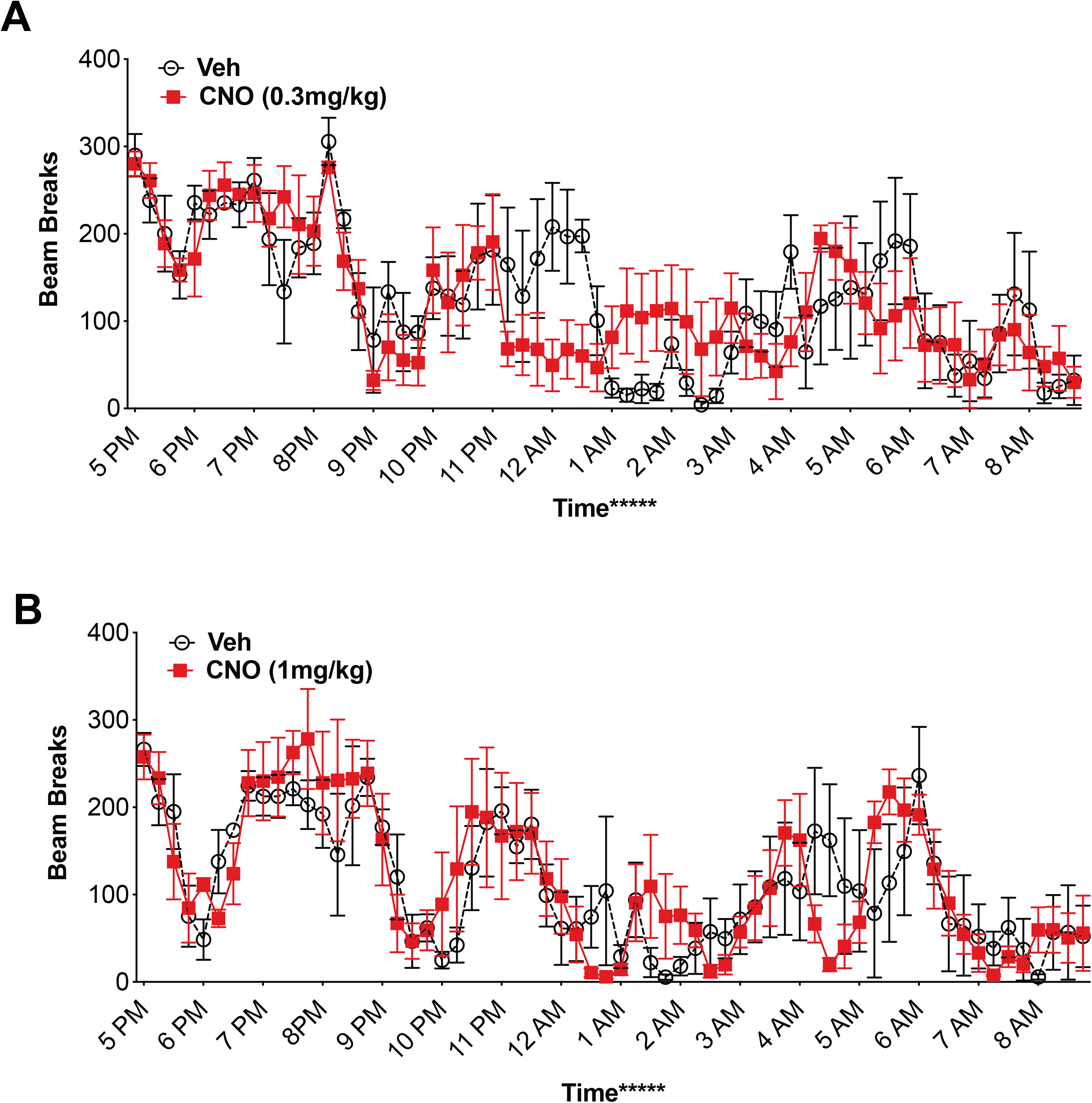
Locomotion after acute injection of Vehicle or CNO (0.3mg/kg or 1mg/kg). **(A,B)** On day one of locomotor behavior testing, mice were given Veh or CNO (0.3 mg/kg **(A)**; 1 mg/kg **(B)**) and immediately placed in the locomotor testing chamber. Data are presented as total beam breaks per 15min-bin. Mean±SEM. n=4/Veh, n=5/CNO. Repeated Measures [RM] Twoway ANOVA, Main effect of Time F_5.562, 38.93_=5.728,***P<0.001, ω_p_^2^=0.35 **(A)**, F_63, 441_=6.996,****P<0.0001, ω_p_^2^=0.37 **(B)**, Main effect of Subject F_7, 441_=5.402,****P<0.0001 **(A)**, F_7, 441_=16.15,**** P<0.0001 **(B)**, Treatment F_1, 7_=0.1478, P>0.05, ω_p_^2^=0.00 **(A)**, F_1, 7_=0.03326, P>0.05, ω_p_^2^=0.00 **(B)**, TimeXTreatment F_63, 441_=1.100, P>0.05, ω_p_^2^=0.01 **(A)**, F_63, 441_=0.9381, P>0.05, ω_p_^2^=-0.01 **(B)**. Veh=vehicle.

**Supplementary Figure 2.**
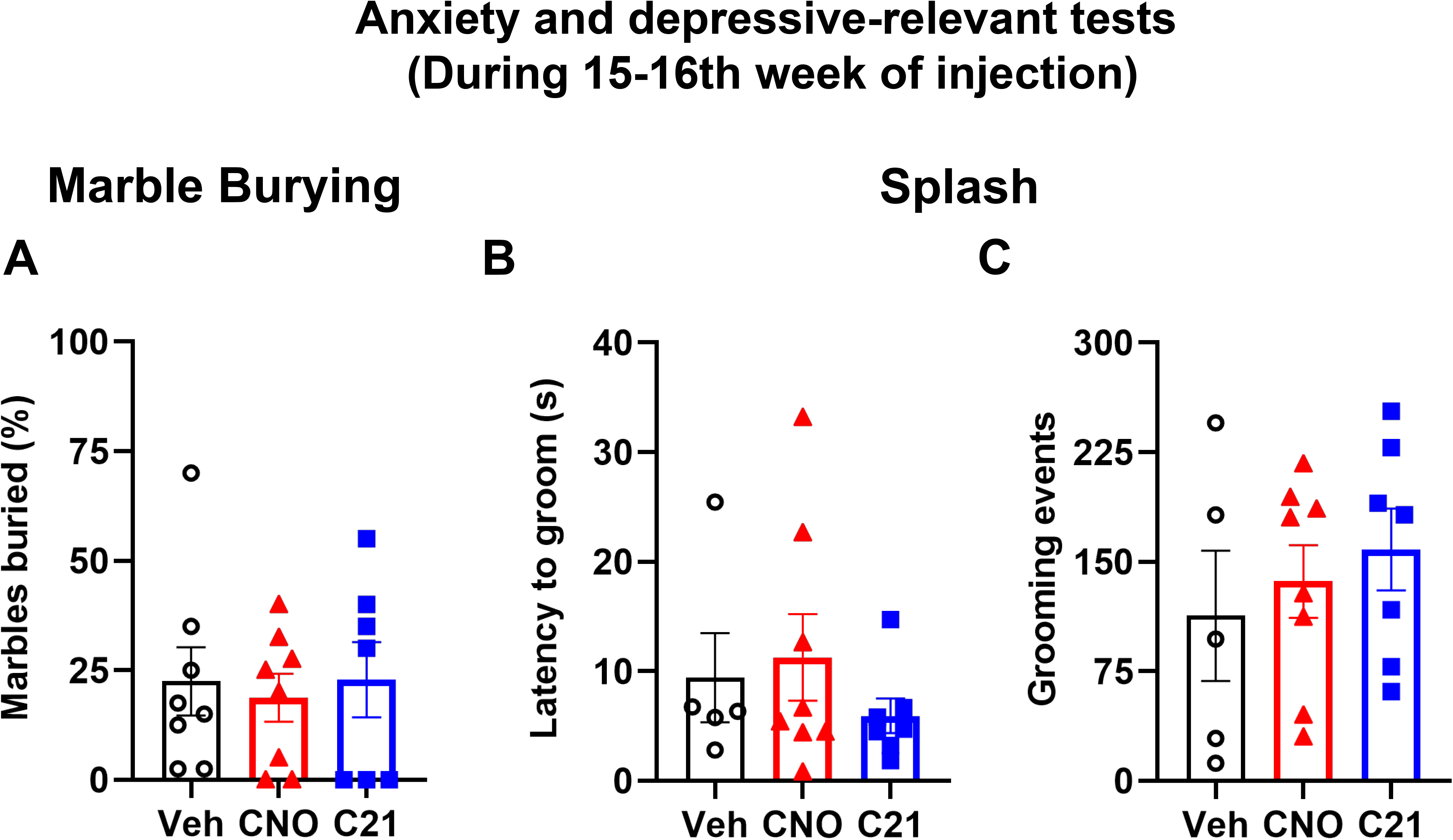
Behaviors relevant to anxiety and depression after fifteen weeks of Veh, CNO, or C21 injections. **(A)** In the marble burying test, mice given Veh, CNO, or C21 buried a similar percentage of marbles. **(B-C)** Due to loss of Veh animals, depressive-like behavior in the sucrose splash-induced grooming test is not conclusive (latency to grooming **(B)**, and grooming events **(C))**. Mean±SEM. Veh n=8 **(A)**, n=5 **(B-C)**; CNO n=8 **(A-C)**; C21 n=7 **(A-C)**. One-way ANOVA, Treatment F_2, 20_=0.09887, P>0.05, ω_p_^2^=-0.09 **(A)**, One-way ANOVA, Treatment F_2, 17_=0.7017, P>0.05, ω_p_^2^=-0.03 **(B)**, One-way ANOVA, Treatment F_2, 17_=0.4806, P>0.05, ω_p_^2^=-0.05 **(C)**.

## Notes

### Competing Interest Statement

The authors have declared no competing interest.

